# Strong cascading impacts of micropollutants on planktonic food web in urban river

**DOI:** 10.1101/2023.12.03.569813

**Authors:** Bob Adyari, Lanping Zhang, Francisco J.A. Nascimento, Yiqing Zhang, Meixian Cao, Jianjun Wang, Hongjun Li, Qian Sun, Changping Yu, Anyi Hu

**Affiliations:** CAS Key Laboratory of Urban pollutant Conversion, Institute of Urban Environment, Chinese Academy of Sciences, Xiamen 361021, China; Fujian Key Laboratory of Watershed Ecology, Institute of Urban Environment, Chinese Academy of Sciences, Xiamen 361021, China; Department of Environmental Engineering, Universitas Pertamina, Jakarta 12220, Indonesia; Department of Ecology, Environment and Plant Sciences, Stockholm University, Stockholm 10691, Sweden; Baltic Sea Centre, Stockholm University, Stockholm 10691, Sweden; University of Chinese Academy of Sciences, Beijing 100049, China; State Key Laboratory of Lake Science and Environment, Nanjing Institute of Geography and Limnology, Chinese Academy of Sciences, Nanjing, 210008, China; State Environmental Protection Key Laboratory of Coastal Ecosystem, National Marine Environmental Monitoring Center, Dalian 116023, P. R. China

**Keywords:** urban river, micropollutants, planktonic food web, urbanization, cascade-effect

## Abstract

Urban rivers rely on intricate multitrophic interactions in food webs, which are vital for ecological functions. Human-induced pollutant discharge, notably micropollutants, poses a threat to these interactions. To study the impact of micropollutants on the microbial food web, we sampled downstream areas of the Jiulong River’s north tributary (lower urbanization, ∼25% built land) and west tributary (higher urbanization, ∼65% built land) for eleven consecutive days in both dry and wet seasons in Fujian, China. We constructed a conceptualized planktonic food web model by employing DNA metabarcoding targeting bacteria, micro-eukaryotes (algae and protozoa), and microzooplankton. Our results revealed that the more urbanized west tributary exhibited significantly higher micropollutant concentrations. Structural equation modeling (SEM) incorporating micropollutants and all food web elements indicated a significantly stronger impact of micropollutants on the planktonic food web in the west tributary than in the north tributary. Micropollutants exhibited both direct and indirect (cascade) effects on high trophic levels (protozoa and microzooplankton), with algal communities mediating the cascade effects instead of bacterial communities, likely due to the increased eutrophication in the urbanized tributary. Overall, our study underscores the potential ecological disruptions caused by micropollutants in shaping planktonic food web interactions in urban rivers.

**Synopsis:** Urban rivers face ecological threats from micropollutants. This study shows elevated micropollutants significantly affected the planktonic food webs. Algal, rather than bacterial, food web pathway mediated micropollutants cascading effects on high trophic levels, emphasizing micropollutant-induced disruptions in urban rivers.

## 1. Introduction

Rivers play a key role in society, such as providing water for domestic and agricultural use, transportation, industrial application, and drinking water. Besides, rivers also carry various important biogeochemical cycles and ecosystem services.^1^ However, the growing trend of urbanization and pollution pose severe problems in the sustainability of the riverine ecosystem.^2^ As a group of pollutants, pharmaceuticals, and personal care products, i.e., micropollutants, have been studied and detected in rivers worldwide in the last two decades.^3^ Micropollutants are known to be pseudo-persistent due to their continuous discharge to the environment.^4^ Despite their occurrences at low concentrations (ngL^-1^ to µgL^-1^) in rivers,^5,6^ micropollutants are designed to interact with a wide range of biological molecules, raising concerns about their deleterious impact on individual and/or disruptive effects at the ecosystem level.^7^ One of the main contributors to micropollutant contamination in rivers is the discharge from the wastewater treatment plant (WWTP) and micropollutant pollution levels have been proportionally associated with the scale of urbanization^5,8^ and bioaccumulation of micropollutants in have been observed in effluent-receiving river. ^6^

Planktonic communities encompassing bacteria, micro-eukaryotes (microscopic algae and protozoa), and microzooplankton are linked by inter-trophic interactions in aquatic food webs. These interactions play a central role in controlling energy and nutrients flow from low to higher trophic levels, sustaining the biogeochemical cycles.^9–11^ Each component of the food web plays a unique role in nutrient cycles. For instance, bacteria play an important role in the decomposition of organic matter and in the recycling of nutrients. They are also important food sources for higher trophic level organisms. Similarly, ciliates and rotifers contribute to nutrient cycling through their feeding and excretion processes, and they also serve as important food sources for organisms at higher trophic levels.^12–14^ Microscopic algae as primary producers are also an important food source for various planktonic predators/grazers.^15^ Multiple studies have documented that each element in the planktonic food web is sensitive to environmental changes such as land-use changes and pollutant exposure^16–18^ and disruption of one element would cause a chain of consequences in the food web and ecosystem functions.^16,19,20^

Pollutants can affect microorganisms through direct exposure to microbial cells,^19^ resulting in various effects at the individual level, such as behavioural impairment, physiological stress, and cell death^4,19^ up to effects at the community level, such as the change in beta diversity^17,21^ and ecosystem functions.^22,23^ To date, many efforts have been made to study the effects of micropollutants on planktonic communities in the field and laboratories.^18,21,23,24^ Our previous study highlighted a strong effect of micropollutant compounds on bacterial communities^18^ but not on micro-eukaryotic communities^24^ in a subtropical river. Separately, algal communities have also been reported to be susceptible to micropollutant exposure in terms of primary productivity^23^ and community composition.^21^ However, most of the previous studies investigating the micropollutant impact on organisms have mostly focused on a single group of organisms such as bacteria^17,18^ or microeukaryotes.^24^

Considering the complex nature of inter-trophic relationship in planktonic communities, the estimated pollutants effect based on a single group of planktonic food web may underestimate the extent of micropollutant impact to the entire food web.^25,26^ Recently, some studies have incorporated simple food web interactions to investigate the effect of pollutants. For example, Rumschlag et al. (2020) reported an indirect effect of pesticide on ecosystem respiration by disrupting zooplankton grazing in algae.^20^ Friman et al. (2015) reported an increased resistance of bacteria to predators under exposure to sublethal antibiotic concentration, resulting in a decrease in predator abundance and diversity.^27^ These lines of evidence have increased our understanding of the effect of micropollutants in a more realistic scenario, although all the aforementioned studies were based on controlled experiments and may not reflect conditions in the field. Therefore, there is a lack of empirical information from field studies with high ecological realism, especially those comparing the effect imposed by high and low levels of micropollutant contamination on the riverine planktonic food web. With the advance of DNA metabarcoding techniques, such as amplicon sequencing, previous laborious efforts to collect samples for studying the multitrophic planktonic food web from the field could be simplified, and large number of samples can be processed in a shorter amount of time.

In this study, the planktonic food web from two tributaries at the downstream of the Jiulong River, China exhibiting different levels of urbanization and micropollutant contamination was compared. Water samples were collected with high temporal resolution (eleven consecutive days) to provide detailed estimates of changes in planktonic communities on a fine time scale,^28,29^ which were recommended to study the rapid response of microorganisms to micropollutants.^30,31^ We used DNA metabarcoding with three different primer sets covering bacteria, micro-eukaryotic (algae and protozoa), and microzooplankton communities representing a simple food web model with bacteria and algae as base food source (i.e., low trophic level),^12^ and protozoa and microzooplankton as consumers (i.e., high trophic level). By using structural equation modelling (SEM), we deciphered how the micropollutant affects each component of the food web to get a comprehensive understanding of the direct and indirect impacts of micropollutants at the food web level. We expected that higher urbanization would result in a higher level of micropollutant contamination, which in turn would have a stronger impact on the riverine planktonic food web. Furthermore, we anticipated both a direct impact of micropollutants on higher trophic levels within the food web and a cascading effect of micropollutants through bacteria or algae.

## 2. Materials and methods

### 2.1. Samples collection and processing

Jiulong River, the second largest river in Fujian Province, China, consists mainly of the north and west tributaries.^32^ At the downstream of this river, the west tributary drainage area encompasses the majority of Zhangzhou city and receives more anthropogenic loads through wastewater treatment plants than the north tributary.^32^ We collected water samples at three sites in the downstream of each tributary for eleven consecutive days in both January and August 2020, representing dry and wet hydrological seasons in Jiulong river,^33^ respectively (**Fig. 1**). The three sampling sites at the downstream of the west tributary had higher built land area than the three sampling sites at the downstream of the north tributary (∼65% vs. ∼25%). Heavy rainfall event was taking place on the 5^th^ day of wet season sampling activity; therefore, sampling was not conducted on that day. This situation also allowed us to observe the temporal (daily) change of riverine abiotic and biotic condition induced by heavy rainfall event. A total of ten micropollutant compounds, which were frequently detected in the investigated area,^34–36^ were determined using a liquid chromatography with triple quadrupole mass spectrometry (LC-MS/MS) as shown in our previous work.^36^

**Fig. 1.**
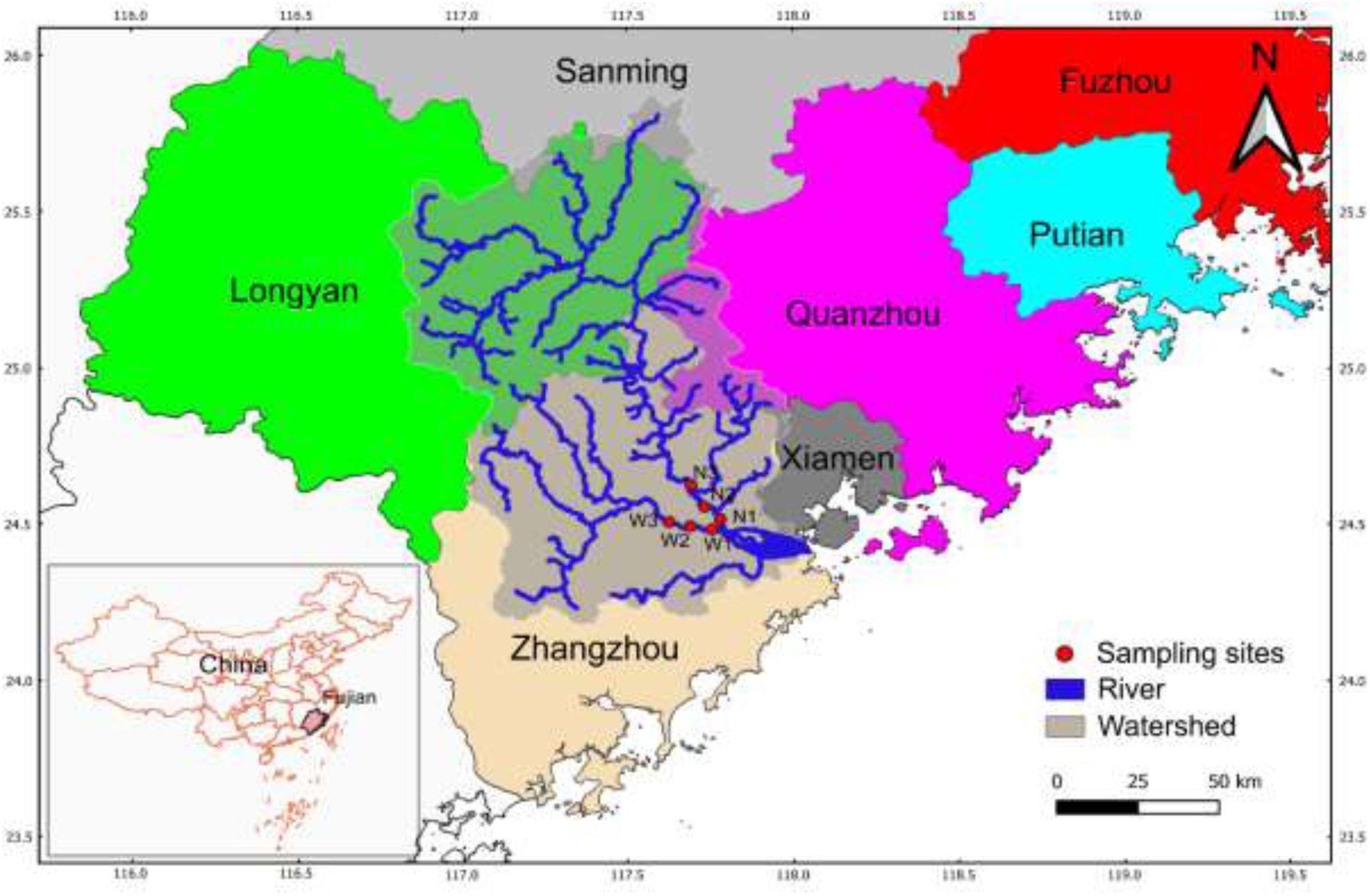
Six sampling sites: three at the downstream of the west tributary (W3, W2, W1) and three at the downstream of the north tributary (N3, N2, N1) of The Jiulong River, Fujian Province, China.

Water samples for DNA analysis (∼1.5 Liter) were filtered through 200 µm nylon mesh, kept in ice box (∼ 4°C), and transported to laboratory. Water samples (∼600 ml) were immediately filtered through 0.2 µm filter (Millipore, Bedford, MA, USA) and stored at −80°C until DNA extraction. Chlorophyl-a was measured using Phyto-PAM phytoplankton analyzer (HeinzWalz, Effeltrich, Germany). Separately, 50 ml of samples were filtered through 0.45 µm filter to determine the concentrations of common inorganic nutrients (NO_2_^-^-N, SO_4_^+^, NO_3_^-^-N, PO_4_^3-^-P, NH_4_^+^-N) and other water ions (F^-^, Cl^-^, Na^+^, K^+^, Ca^2+^, and Mg^2+^) using ion chromatograph system (DionexICS-3000 and Dionex Acquion, Thermo Fisher, USA). Built land area percentage within 2 km radius of each site was determined based on Landsat-7 ETM+ (2020) data with a resolution of 30 m with QuantumGIS.^37^ This built land area was used as proxy for the urbanization rate.

### 2.2. DNA extraction and metabarcoding

DNA from a total 132 samples were extracted using FastDNA SPIN kit for Soil (Qbiogene-MP Biomedicals, Irvine, CA, USA) according the manufacturer protocol. The bacterial community, micro-eukaryotic community, and microzooplankton community data were obtained by amplifying the hypervariable V4-V5 region of bacterial 16S rRNA genes, V9 region of eukaryotic 18S rRNA gene, and a region of cytochrome c oxidase subunit I (COI), respectively. The details of primers and bioinformatics processing is described in **Supplementary Information S1**. Before further analyses, for each sample, the number of reads for bacteria, micro-eukaryotes, and microzooplankton were rarefied to 37,881, 26,046, and 3,000 reads respectively. Samples with too low read numbers were discarded which resulted in the total of 124 samples (dry season west tributary: 29, dry season north tributary: 31, wet season west tributary: 32, wet season north tributary: 32).

### 2.3. Statistical analysis

Prior to model the planktonic food web relationships, we classified the microeukaryotes into two functional guilds indicative of their lifestyle^38^: photosynthetic micro-eukaryotes (hereafter, algae) comprising Ochrophyta, Chlorophyta, and Cryotphyta and consumer micro-eukaryotes (hereafter, protozoa) comprising Ciliophora and Opalozoa.

Principal coordinate analysis (PCoA) based on Bray-Curtis distance was used to visualize the spatio-temporal (i.e., sampling sites vs daily change within eleven days sampling campaign) pattern of the riverine planktonic food web elements (bacteria, algae, protozoa, and microzooplankton). Permutational multivariate analysis of variance (Permanova) based on the same distance was used to calculate the variance explained by spatial and temporal factors. Subsequently, micropollutants, physico-chemicals, bacteria, algae, protozoa, and microzooplankton data were integrated with Diablo (data Integration Analysis for Biomarker discovery using Latent cOmponents) through *block.splsda* function from the Mixomics R package.^39^ Diablo is a framework with the purpose of integrating multiple data sets in omics study and identifying the latent components that explain the variations in the data.^17^ Prior to analyses, only bacterial and micro-eukaryotic ASVs with an occurrence (minimum one sequencing read) in at least 50% of the samples, and microzooplankton ASVs with an occurrence in at least 40% of the samples were included to the analyses to reduce the noise of the data.^40^ To account for the differences in sequencing depths among bacteria, micro-eukaryotes and microzooplankton, data were transformed using total sum scaling and centered log-ratio for each domain.^40^ Physico-chemicals and micropollutant variables were log10(x+1) transformed to improve linearity. Diablo integration produces latent components representing the main gradient of changes for each biotic and abiotic groups which was used further for SEM analysis (**Fig. S1**).

SEM was utilized to understand how micropollutants and physico-chemicals impact the riverine planktonic food web. It involved components like bacteria, algae, protozoa, and microzooplankton in the food web, along with physico-chemicals and micropollutants as abiotic factors. Urbanization’s spatial aspect (measured as built land percentage) and temporal changes during daily sample collection were also considered. Before conducting SEM, a hypothesized model (**Fig. S1**) was developed, depicting a bottom-up scenario where micropollutants affect the food web through bacteria and algae, which serve as food sources for higher trophic level organisms like protozoa and microzooplankton.^12^ Microzooplankton were assumed to primarily feed on algae and protozoa.^41,42^ The built land percentage and daily change were proxies for the spatio-temporal patterns^43,44^ explaining the changes in the abiotic and food web. The SEM, constructed using the ‘psem’ function of the PiecewiseSEM package, accounted for non-independence of sampling sites using a linear mixed effect model.^45,46^

SEM is a powerful tool for uncovering direct and indirect causes within complex models,^47^ allowing the tracing of how micropollutants impact consumers through their food sources, like bacteria and algae. It partitions total effects into direct and indirect impacts, enabling a clearer understanding of the pathways through which influences propagate.^45–48^ For example, micropollutants can directly affect protozoa communities (direct effect). Additionally, they can indirectly impact protozoa by first affecting bacteria and algae communities. Bacteria and algae act as mediators in the way micropollutants influence protozoa. This separation of indirect effects helps determine the specific pathways, such as whether micropollutants affect protozoa through bacteria or algae in the food web. The direct, indirect, mediator, and total effects were standardized and their significances were calculated based on 1000 permutations through bootEff function of semEff package^47^. The multicollinearities among predictors were checked with RVIF function from semEff package^47^. We used Wilcoxon-test to test the differences between the physico-chemicals’s and micropollutants’s total effects on the food web (bacteria, algae, protozoa, microzooplankton) in the west and north tributaries of Jiulong River.

## 3. Results

### 3.1. Higher urbanized area and higher micropollutants in the west than the north tributaries despite high similarity taxonomy composition at the phylum/Division level

The geospatial land-use analysis indicated that the average built land area from the three sites in west tributary (average 65%) was ∼3 times higher than that of north tributary (27.2%) (**Fig. S2**). Likewise, the mean concentration of total micropollutants was also significantly higher in west than north tributaries in dry (390.13 ng/L^-1^ vs. 86 ng/L^-1^, *p* < 0.05) and in wet season (115.85 ng/L^-1^ vs. 73.82 ng/L^-1^, *p* < 0.05) (**Fig. S2**). Despite, the difference in the built land area percentage and micropollutants concentration, the taxonomy composition of planktonic food web in the north and west tributaries at phylum/division level were similar. The bacterial communities at the phylum level were dominated by Proteobacteria, Bacteroidota, Actinobacteria in west and north tributary (**Fig. S3a**). Cyanobacteria appeared as one of the most abundant phyla only during wet season in both tributaries and decrease sharply in the dry season. The major phyla of microeukaryotes belong to that of photosynthetic group comprising Chlorophyta, Cryptophyta, and Ochrophyta that contributes 66.8% – 71.2% of north tributary and 72% – 74.2% of west tributary micro-eukaryotic communities (**Fig. S3b**). On the other hand, non-photosynthetic micro-eukaryotic group was dominated by consumer such as Ciliophora and Opalozoa which accounted for 16.6 – 18.9% of north tributary and 10.9 – 14.9% of west tributary micro-eukaryotic communities. For the subsequent analysis, we focus specifically to these two groups as they represent groups with most definite lifestyle (i.e., primary producers and consumers respectively). Microzooplankton communities in both tributaries were mainly dominated by rotifera and arthropoda which accounted for 94.1 – 98.6% of north tributary and 86.6 – 94.5% of west tributary microzooplankton communities (**Fig. S3c**).

### 3.2. Different spatio-temporal pattern of the riverine planktonic food web communities between the north and west tributaries

Spatial pattern (differences among sampling sites in the same tributaries) appeared to be stronger for bacterial, algae, protozoa, and microzooplankton communities in north (R^2^ = 0.23 – 0.59) than west (R^2^ = 0.11 – 0.48) tributary, while daily variation was a more important factor for the planktonic food web communities in the west (R^2^ = 0.08 – 0.21) than north tributary (R^2^ = 0.03 – 0.07) (**Fig. 2**). A heavy rainfall event occurred during the wet season sampling campaign (proxied by temporal daily change) played a remarkable role in shaping the food web communities in west tributary (R^2^ = 0.08 – 0.21), while such impact was less pronounced in the north tributary (R^2^ = 0.03 – 0.07). Furthermore, regardless of tributary or season, spatial variations of the low trophic groups (i.e., bacteria or algae) (R^2^ = 0.4 – 0.59) were consistently higher than that of consumers such as protozoa and microzooplankton (R^2^ = 0.23 – 0.48) (**Fig. 2q**).

**Fig. 2.**
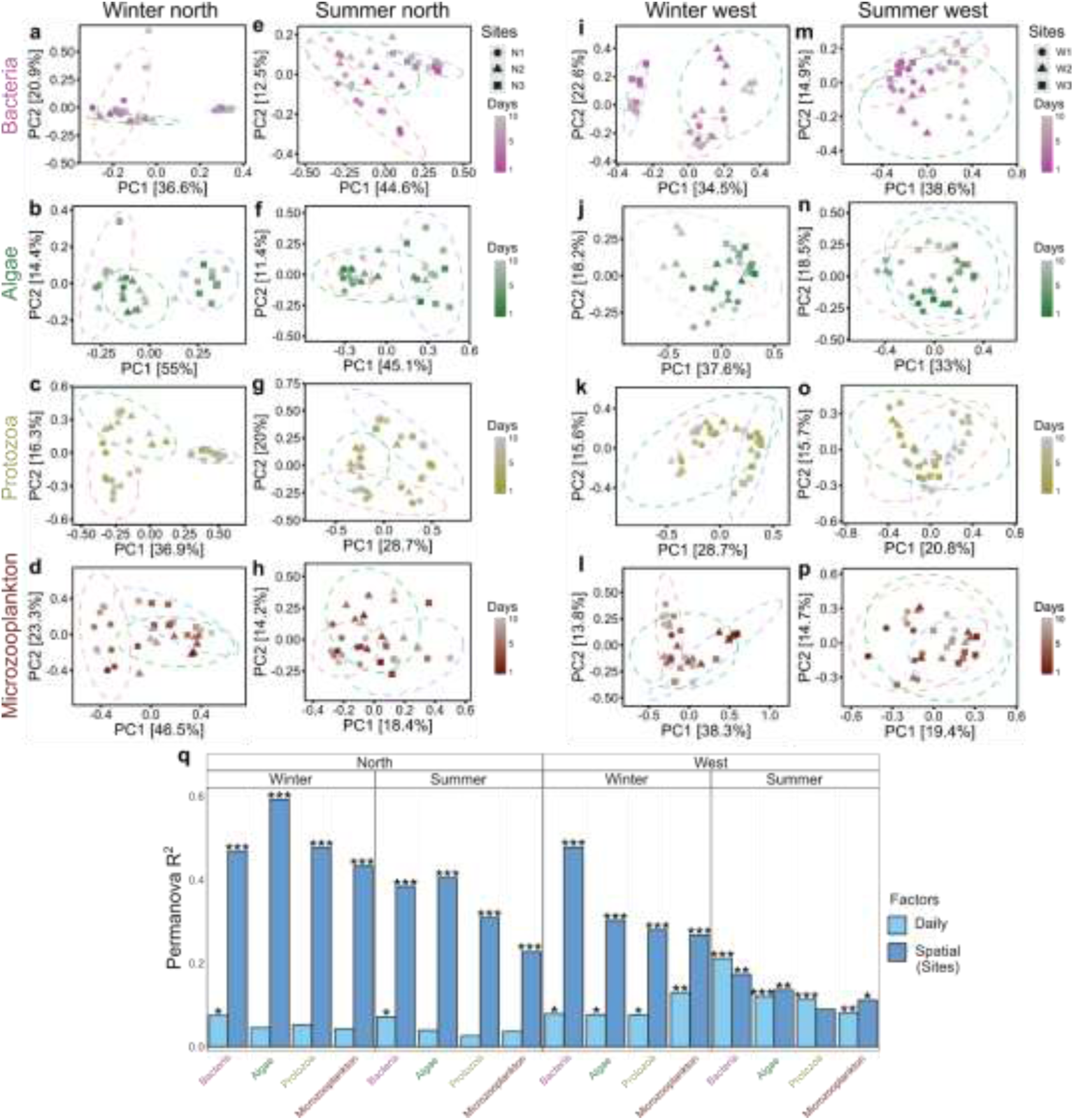
**a-p,** Principal coordinate analysis (PCoA) of bacteria, algae, protozoa, and microzooplankton at ASV levels and **q**, their amount of variation explained (R^2^) by daily and spatial factors from Permanova test for north and west tributaries of The Jiulong Rivers in winter and summer seasons (\*\*\**p* < 0.001; \*\**p* < 0.01; \**p* < 0.05).

### 3.3. Strong link between urbanization and micropollutants impact on planktonic riverine food web

We built four SEM models to investigate the interplay amongst built land area change, daily change (i.e., temporal variation of daily sampling), physico-chemicals, micropollutants, and planktonic food web communities in the north and west tributary of the Jiulong River in dry and wet season seasons. SEM results showed significant direct effects of micropollutants on dry season bacteria (0.24), algae (0.4), and protozoa (0.15) and on wet season bacteria (0.21) and algae (0.33) in the west tributary, while no significant direct effect of micropollutants on any of the food web element was observed in the SEM results for the north tributary in both wet and dry season (**Fig. 3**). It is also worth to note that, the direct effects of micropollutants on the planktonic food web had a decrease trend from low trophic levels, that is, algae (0.33 – 0.4) and bacteria (0.21 – 0.24), to consumers including protozoa (0.1 – 0.15) and microzooplankton (−0.01 – 0.1), indicating that the impact of micropollutants direct exposure tended to decrease towards higher trophic level communities at the expense of their indirect effects (i.e., effects on protozoa and microzooplankton that is mediated by bacteria and algae). For instance, the indirect effect of micropollutants occupied the major proportion in micropollutants total effect on microzooplankton (direct = −0.01, indirect = 0.12) (**Fig. 4a**). In contrast, both the direct and indirect effects of micropollutants were equally strong and significant in protozoa (direct = 0.1, indirect = 0.09) and microzooplankton (direct = 0.1, indirect = 0.06) in the west tributary during wet season.

**Fig. 3.**
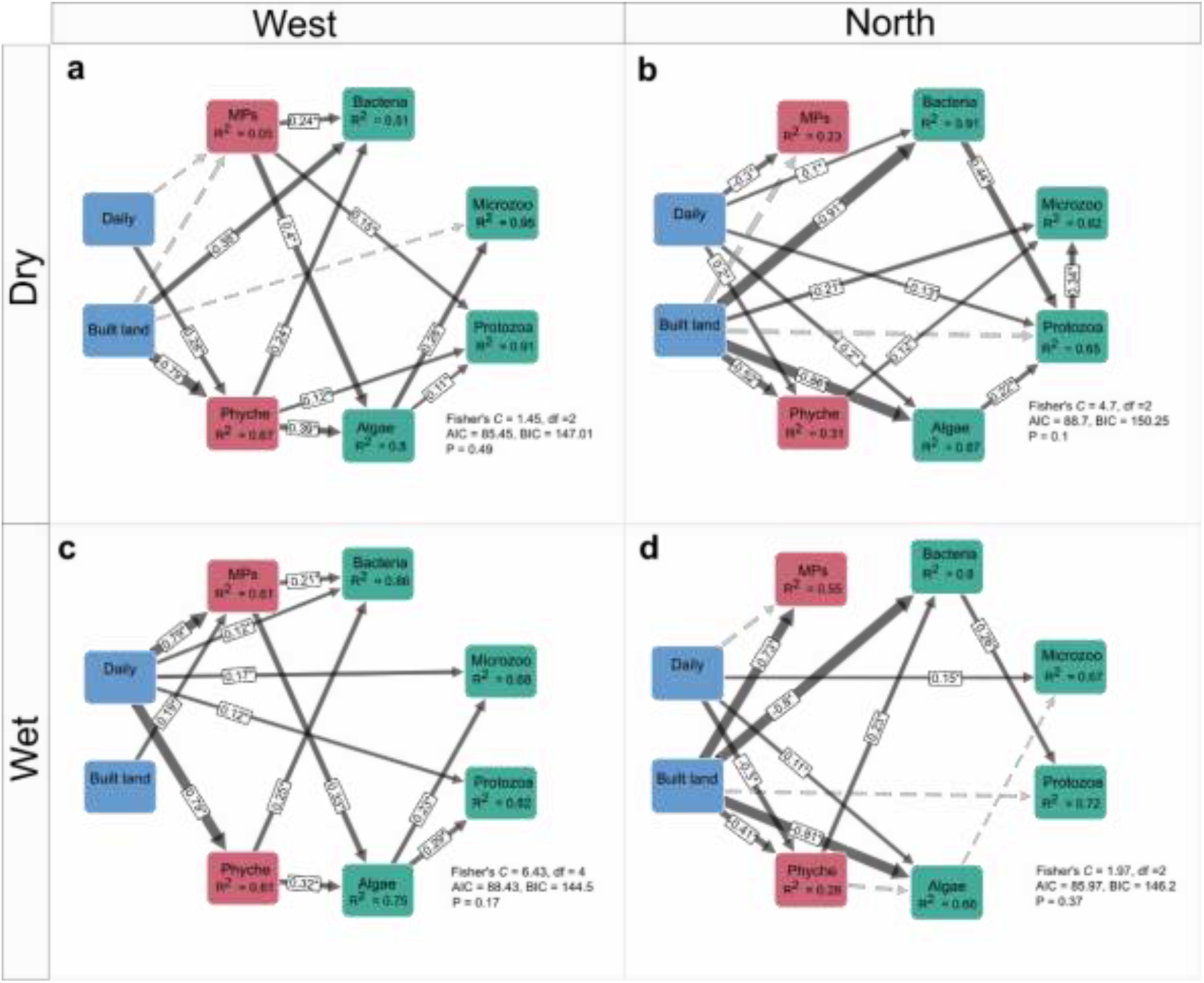
**a-d,** Structural equation model (SEM) based on PiecewiseSEM approach investigating the relationships among daily variation (**Daily**), built land (**Built land**), physico-chemicals (**Phyche**), micropollutants (**MPs**), bacteria (**Bacteria**), micro-eukaryotic algae (**Algae**), protozoa (**Protozoa**), and microzooplankton (**Microzooplankton**). Solid and dashed paths represent significant and insignificant standardized direct effects respectively. Path sizes are proportional to the strength of the coefficients and only arrows with coefficients > 0.1 are displayed for a clear visual representation. The significance of all path coefficients is obtained from 1,000 bootstraps (*p* < 0.05).

**Fig. 4.**
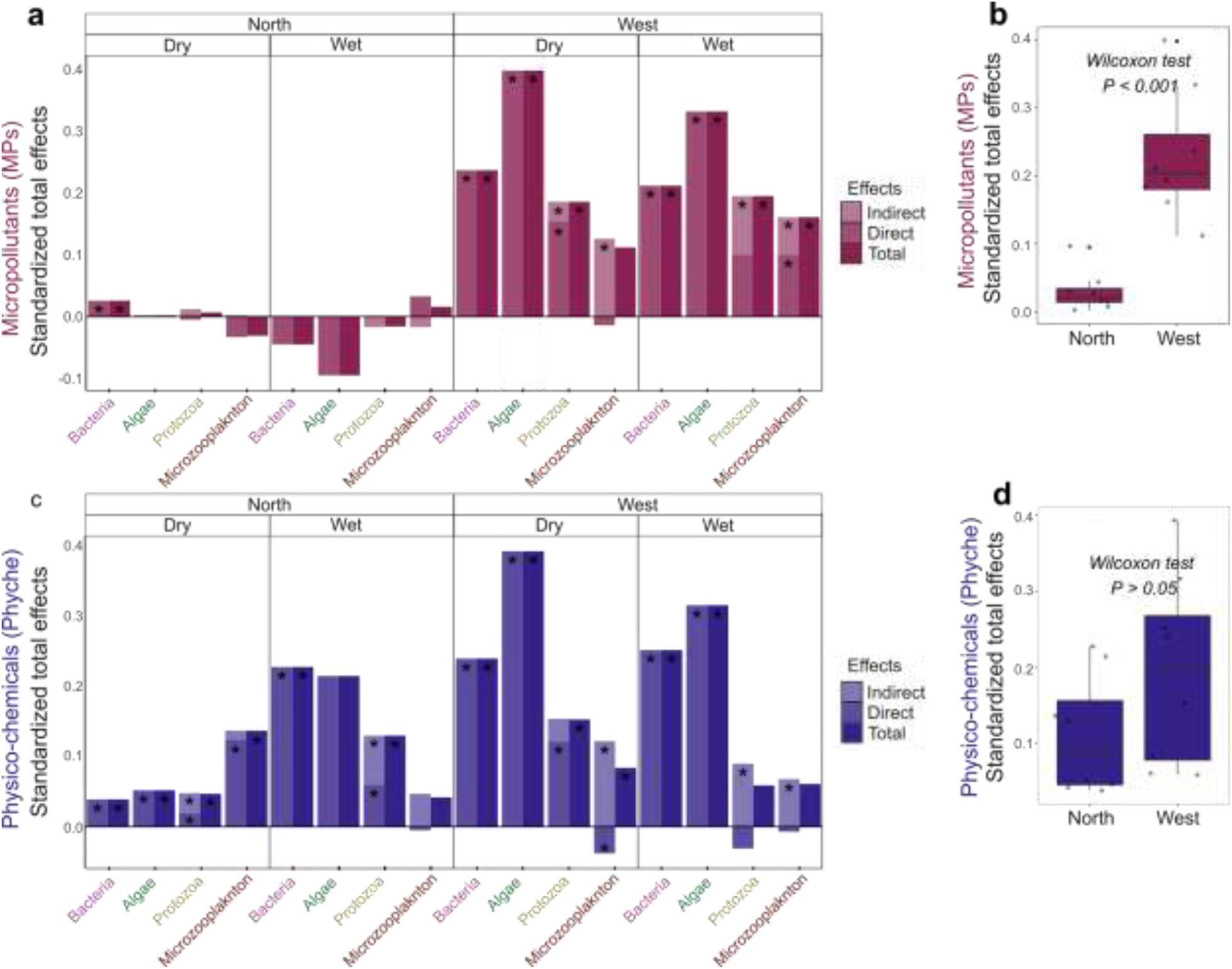
**a,** Standardized total effects of MPs on microbial food web (bacteria, algae, protozoa, and microzooplankton). **b,** Comparison of the standardized total effects of micropollutants on microbial food web between west and north Tributaries of The Jiulong Rivers. **c**, Standardized total effects of physico-chemicals on microbial food web **d,** Comparison of the standardized total effects of physico-chemicals on microbial food web between west and north tributaries of The Jiulong Rivers. The significance of direct, indirect, and total effects are obtained from 1,000 bootstraps (*p* < 0.05).

Similarly, the direct effects of physico-chemical variables on the riverine planktonic food web were more prominent in west than north tributary (**Fig. 3**). However, there was no significant difference (*p* > 0.05) between the total effects of physico-chemicals in north and west tributary (**Fig. 4d**). Contrary, the total effect of micropollutants on food web were significantly higher in west than north tributaries (**Fig. 4b**), implying the importance of micropollutants as an important factor governing the planktonic food web in highly urbanized tributary. Additionally, the impact of micropollutants and physico-chemical variables were equally strong (*p* > 0.05) on the food web in the west tributary but the impact of micropollutants was significantly weaker than physico-chemicals in the north tributary (*p* < 0.001) (**Fig. S4**).

### 3.4. Algae, rather than bacteria mediated the micropollutant impact to high trophic levels through interactions in planktonic riverine food webs

By separating the base of our food web model to bacterial and algal communities, SEM can be used to investigate the pathway of the indirect (cascade) effect of micropollutants and physico-chemicals to consumers (i.e., cascade effect through bacteria and algae). SEM results suggested that algae were the main significant mediator of the cascade effects from micropollutants to protozoa (0.043 – 0.95) and microzooplankton (−0.015 – 0.115) in the west tributary (**Fig. 5a**), while bacteria were more pronounced as mediator of micropollutants for protozoa (−0.012 – 0.011) in north tributary despite the low impact of micropollutants in the north tributary. Similarly, algae were also the significant mediator of physico-chemicals to protozoa (0.042 – 0.09) and microzooplankton (0.065 – 0.113) in the west tributary, while bacteria, algae, and protozoa were all significant mediators in the north tributary (**Fig. 5b**). This results altogether suggested a more diverse potential mediators for cascade impact of micropollutants in the north than the west tributary despite their lower effects to high trophic communities.

**Fig. 5.**
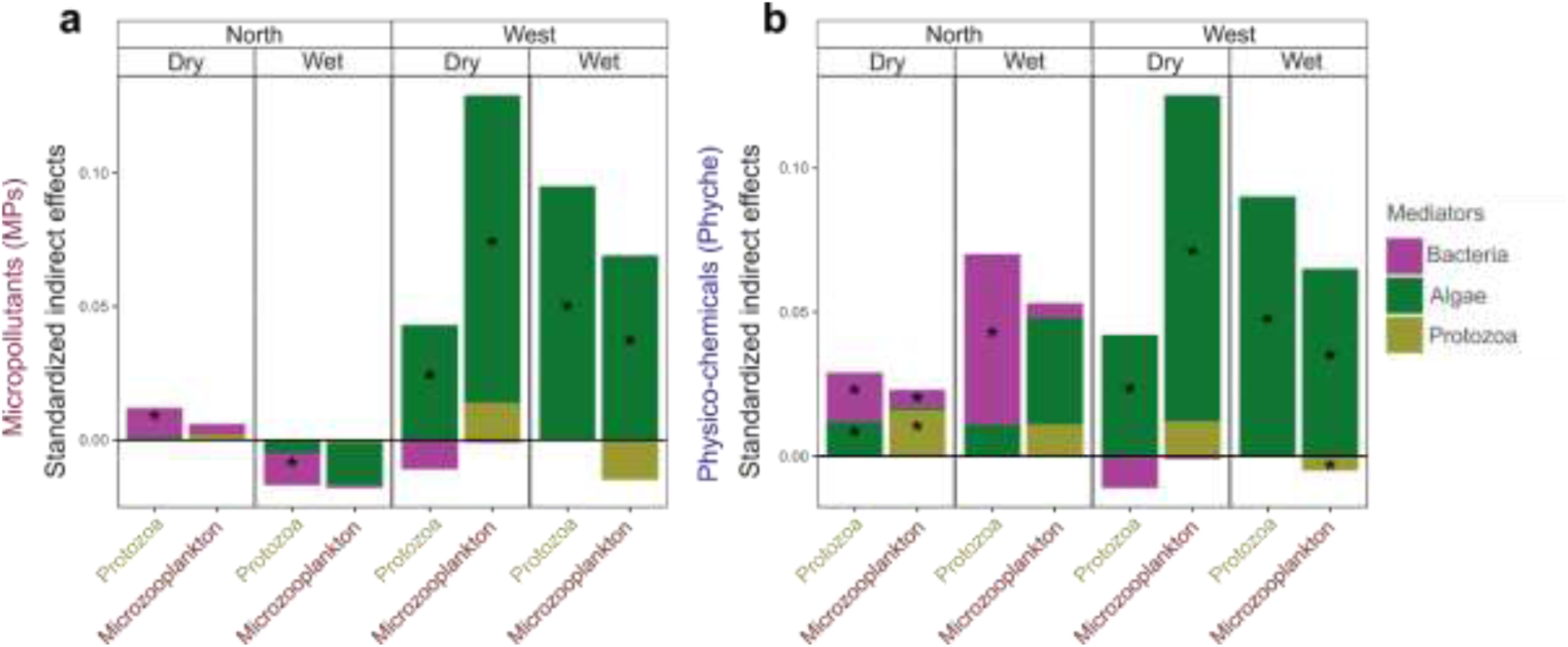
**a,** The proportion of bacteria, algae, and protozoa as mediators for the indirect effect of micropollutants on protozoa and microzooplankton. **b**, The proportion of bacteria, algae, and protozoa as mediators for the effect of physico-chemicals on protozoa and microzooplankton. The significance of the effects is obtained from 1,000 bootstraps (*p* < 0.05).

## 4. Discussion

Urbanization has substantially heightened anthropogenic contaminants in riverine ecosystems, specifically linking increased micropollutant levels with urbanization pressure.^5,8,49^ Notably, highly urbanized downstream river basins pose a more severe threat to estuaries and coastal areas, given their proximity and limited water recovery time.^1^ In our research, the west tributary, characterized by a higher percentage of built-up areas, exhibited significantly elevated daily micropollutant concentrations (**Fig. S2**). Previous studies have emphasized that urbanization significantly influences micropollutant concentrations in surface water, with instances like the Huangpu River in Shanghai experiencing up to a sixfold increase from rural to urban segments.^49^ In line with our initial hypotheses, the impact of micropollutants on the riverine planktonic food web in the west tributary was markedly stronger compared to the north tributary. Additionally, in the west tributary, the force exerted by micropollutants was equally influential as physico-chemical variables (**Fig. S4**), aligning with findings in Mediterranean rivers that suggested a robust joint effect of these factors on biofilm and invertebrate communities.^50^

It’s essential to note that factors controlling micropollutants appeared to vary with seasons (**Fig. 3**). For instance, in the wet season, the built land percentage along the sampling sites in the north tributary explained micropollutant variation, while temporal changes were the primary predictor in the west tributary (**Fig. 3**). Heavy rainfall significantly impacted water quality parameters^51^, including micropollutants, as indicated by the explanatory power of temporal changes in our SEM results (**Fig. 3c**). Previously, Corcoll et al. (2014) also suggested that water flow was an important factor mediating the effect of micropollutants on freshwater biofilm.^52^ Our study indicates that such effects may impact not only biofilms, but also other components of the planktonic food web. Heavy rainfall could exert a strong pulse and increase water volume, while also washing land pollutant, causing leachate to leak into the open environment, which could facilitate the introduction of micropollutants into the water body.^53^ Despite a significant influence of built land area on micropollutants in the north tributary, their impact on the planktonic food web was not noteworthy. In fact, it was notably lower than the impact observed in the west tributary. Previous studies have shown a relatively lower impact of micropollutants on riverine communities compared to physico-chemical variables. For instance, Adyari et al. (2020) found physico-chemical variables exerted a stronger influence on shaping bacteria communities in a deep water reservoir than micropollutants.^17^ Similarly, Sabater et al. (2016) reported a stronger influence of physico-chemical and land use factors compared to micropollutants on riverine biofilms and invertebrate communities.^50^ Micropollutants are known to be toxic for some bacterial species while serving as a substrate for others. ^10,54^ Therefore, the lower impact of micropollutants on the planktonic food web in the north tributary may be attributed to their low concentration, possibly insufficient to trigger responses from bacterial species.^55^

Our findings reveal that the cascade effects of micropollutants on higher trophic level biota in the planktonic food web, such as protozoa and microzooplankton, are predominantly mediated by algae rather than bacteria (**Fig. 5**). We propose that the elevated concentration of nutrients and dissolved organic matter,^51^ characteristic of eutrophic ecosystems in the west tributary, creates a more favorable environment for algae growth. This is supported by significantly higher chlorophyll-a concentration (**Fig. S2***, p* < 0.05) in the west tributary, indicating higher eutrophication levels and algal biomass compared to the north tributary.

Eutrophication plays a crucial role in determining the significance of bacteria and algae as food sources for higher trophic levels in aquatic ecosystems.^42,56^ It is hypothesized that bacteria play a more critical role as a basal resource in nutrient-poor environments, while high primary production in nutrient-rich environments makes algal communities a more abundant food source.^13,42,56^ Additionally, bacteria are considered a low-nutrient food source, acting as a supplement to algae. ^42^ Exposure of micropollutants in algae communities can lead to various adverse effects, such as growth inhibition,^57^ decreased primary productivity,^4^ and changes in community structure,^21,24^ ultimately affecting protozoa and microzooplankton through alterations in consumption patterns.^27^ Guacsh et al. (2016) reported that Diatom (algae) communities have been reported to be more susceptible to grazing pressure in environmentally relevant micropollutant exposure treatments than under normal conditions.^16^ Furthermore, algae cells have the capability to adsorb and degrade micropollutant compounds,^58^ potentially leading to the accumulation of micropollutants in higher organisms through the food web.^59^ While the specific mechanisms of how algae mediate the impact of micropollutants to higher trophic levels are complex, our study provides empirical evidence that algae play a significant and crucial role in the urban river planktonic food web, with a high potential for cascading the impact of micropollutants within the food web.

Previous experimental studies reported that multiple micropollutants affected more than one trophic level biotic component of the aquatic ecosystem.^60,61^ Despite this, very little field study has been conducted to observe such a phenomenon. To our knowledge, this is the first field study to empirically compare the effect of micropollutants between two water bodies with different levels of urbanization and micropollutants, which is a clear missing piece of evidence to prove that urbanization can lead to a stronger influence of micropollutants on the planktonic food web (**Fig. 6**).

**Fig. 6.**
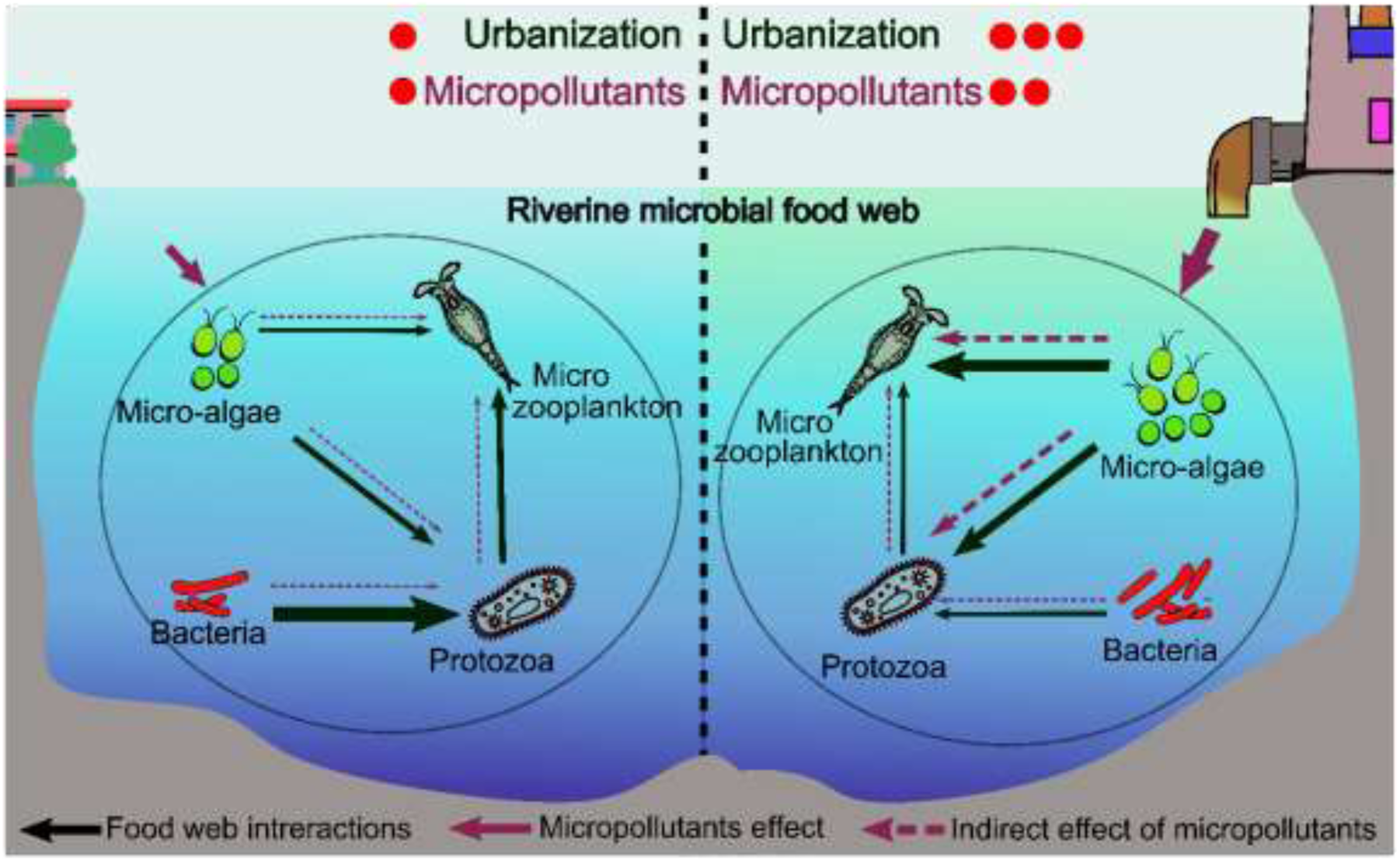
Conceptual diagram of the influence of urbanization and micropollutants on riverine planktonic food web. The width of arrows indicates the strength of relationships

The inclusion of multiple trophic levels enabled us to distinguish the relative importance of different low trophic levels (bacteria or algae) and micropollutants on higher trophic level consumers (protozoa and microzooplankton). Our SEM results suggested that the low trophic levels were also important factors in controlling the higher trophic level consumers (protozoa and microzooplankton) in the planktonic food web in the north and west tributaries (**Fig. S5**). This finding highlights that the non-negligible role of biotic interaction (i.e., inter-trophic relationships) in assessing the impacts of physico-chemical variables or micropollutants factors on the food web in realistic ecosystem scenario.^62^ In summary, our field study contributes novel insights into the intricate relationship between urbanization, micropollutants, and riverine planktonic food webs. It highlights the need for integrated management approaches to mitigate the impact of urbanization on river ecosystems and underscores the importance of considering multiple trophic communities when evaluating the effects of micropollutants.

## Associated content

### Supplementary information

Primers and bioinformatics processing producures, flowchart of data analysis and SEM hypothetical diagram, comparison of built land key percentage, physico-chemical parameters, and micropollutants concentrations between the north and the west tributaries, taxonomy composition (Bacteria, micro-eukaryotic, and microzooplankton) in the north and west tributaries, comparison of the total effect of micropollutants and physico – chemicals effect on planktonic food web in the north and the west tributaries (SEM), the proportion of micropollutants, physico-chemicals, algae, and protozoa, to the total effects of bacteria, algae, protozoa, and microzooplankton (SEM).

## Notes

The authors declare no competing financial interest.

## Acknowledgement

We thank Dr. Dandan Izabel-Shen for valuable comments and discussion on early version of this manuscript. This work was supported by the National Natural Science Foundation of China (31870475 and U1805244). BA was supported by the CAS-TWAS President’s scholarship for international students.

