## supplementary materials for "Strong cascading impacts of micropollutants on planktonic food web in urban river"

**Supplementary Information S1**

The V4-V5 region of 16S rRNA gene was amplified with universal primers 515F (5'-GTG YCA GCM GCC GCG GTA-3') and 907R (5'-CCG YCA ATT YMT TTR AGT TT-3') to profile bacterial community data. The hypervariable V9 region of 18S rRNA gene was amplified with 1380F (5′-CCC TGC CHT TTG TAC ACA C-3′) and 1510R (5′-CCT TCY GCA GGT TCA CCT AC-3′) to profile micro-eukaryotic community data. The  region of cytochrome c oxidase subunit I (COI) was amplified  with mlCOIintF (5'-GGW ACW GGW TGA ACW GTW TAY CCY CC-3′) and mlCOIintR (5' GGR GGR TAS ACS GTT CAS CCS GTS CC-3′) to profile microzooplankton community.[1] Bacterial and micro-eukaryotic raw paired-end sequences were processed and clustered into amplicon sequencing variant (ASVs) with DADA2 pipeline[2] as described elsewhere (https://benjjneb.github.io/dada2/tutorial.html). Subsequently, the taxonomy assignment for bacteria and micro-eukaryotic was performed using RDP classifier with SILVA v132[3] and PR2[4] databases, respectively. Microzooplankton paired-end raw sequences were processed and clustered into ASVs with MetaWorks v1.9.0,[5] and taxonomy assignment was performed with CO1 Classifier v4.0.1.[6]

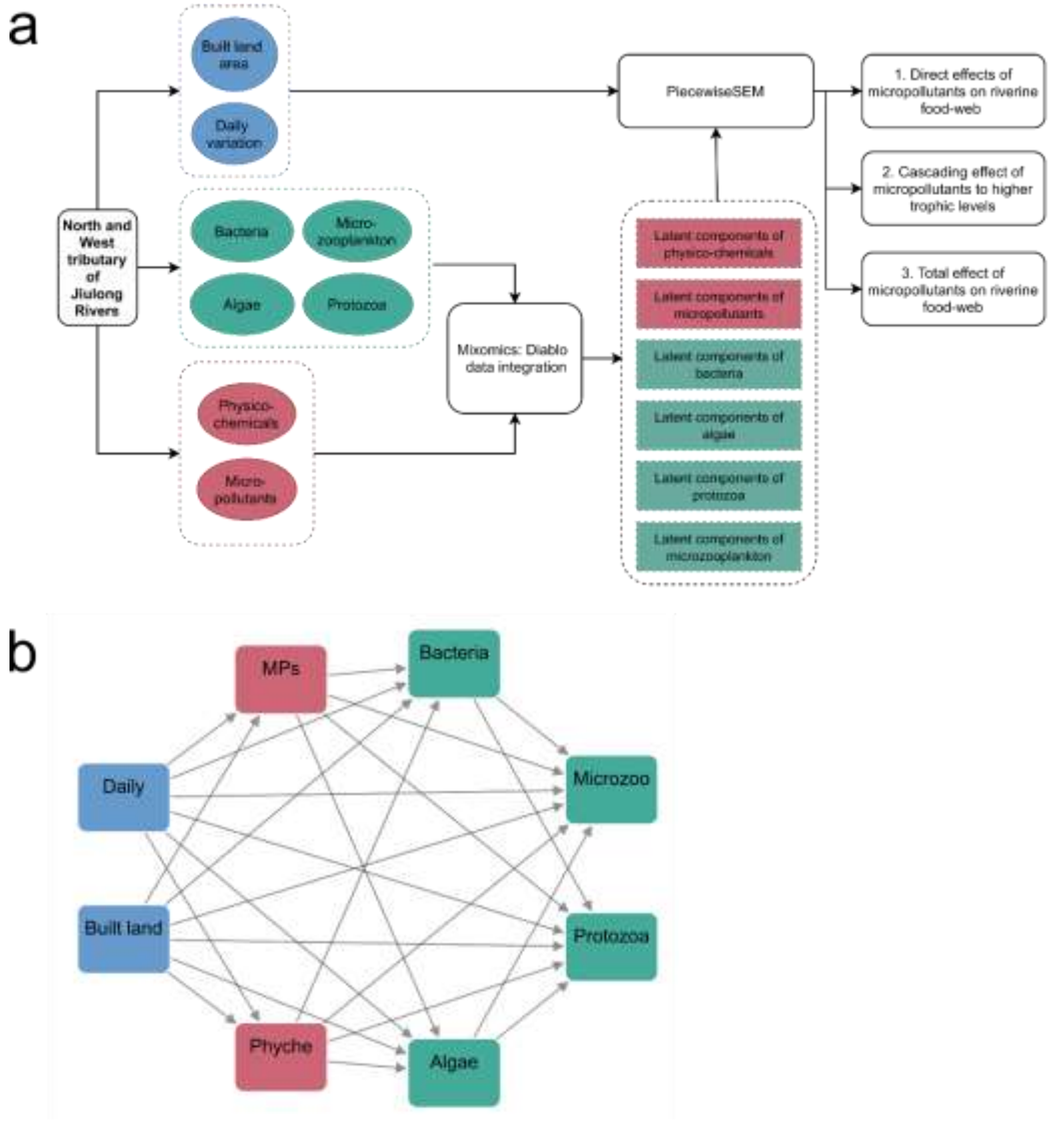


**Fig. S1. a**, Flowchart of integrated analyses with Diablo and SEM. **b**, hypothetical SEM diagram

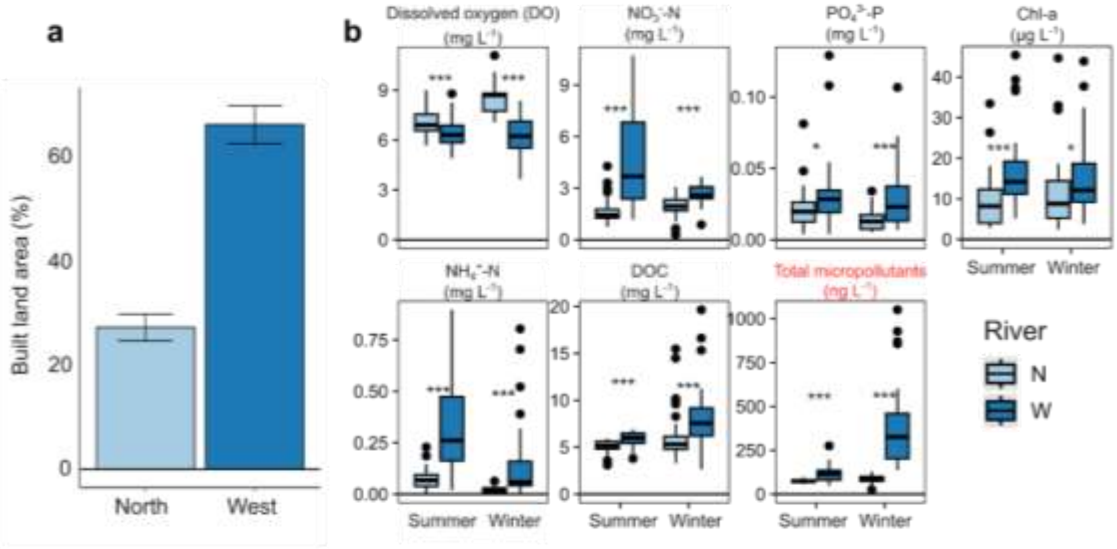


**Fig.S2. a,** Comparison of built land area percentage and **b**, comparison of key physico-chemicals parameters and total micropollutants between the North and West tributaries of The Jiulong Rivers. The significance of the differences is tested using the Wilcoxon test (***$p$ < 0.001; **$p$ < 0.01; *$p$ < 0.05).

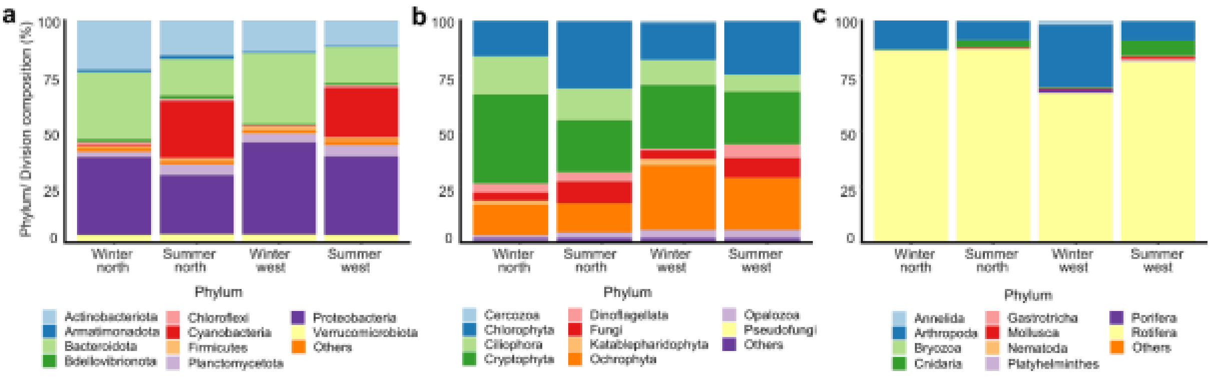


**Fig. S3.** Taxonomy composition of bacteria (**a**), microeukaryotic (**b**), and microzooplankton (**c**) communities in the North and West tributaries of The Jiulong Rivers.

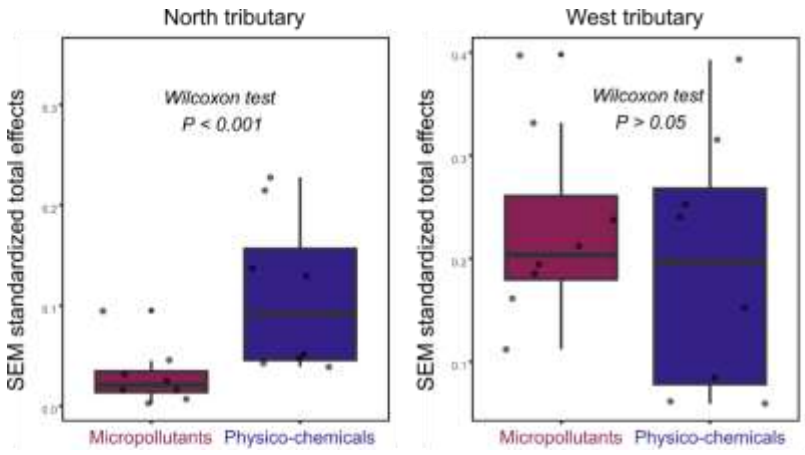


**Fig. S4.** The comparison of the total effect of micropollutants and physico-chemicals effect on planktonic food web in the North and West tributary of The Jiulong River.

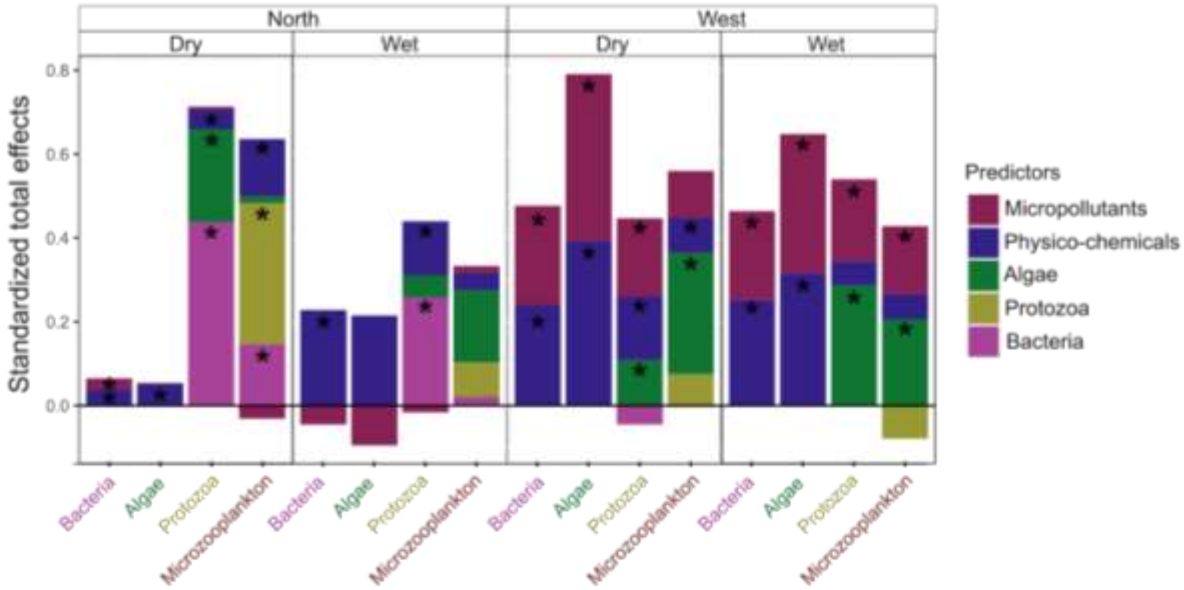


**Fig. S5.** The proportion of micropollutants, physico-chemicals, algae, and protozoa, to the total effects of bacteria, algae, protozoa, and microzooplankton. The significance of total effects is obtained from 1,000 bootstraps ($p < 0.05$).
